# Learnt representations of proteins can be used for accurate prediction of small molecule binding sites on experimentally determined and predicted protein structures

**DOI:** 10.1101/2023.09.07.556685

**Authors:** Anna Carbery, Martin Buttenschoen, Rachael Skyner, Frank von Delft, Charlotte M. Deane

## Abstract

Protein-ligand binding site prediction is a useful tool for understanding the functional behaviour and potential drug-target interactions of a novel protein of interest. However, most binding site prediction methods are tested by providing crystallised ligand-bound (holo) structures as input. This testing regime is insufficient to understand the performance on novel protein targets where experimental structures are not available. An alternative option is to provide computationally predicted protein structures, but this is not commonly tested. However, due to the training data used, computationally-predicted protein structures tend to be extremely accurate, and are often biased toward a holo conformation. In this study we describe and benchmark IF-SitePred, a protein-ligand binding site prediction method which is based on the labelling of ESM-IF1 protein language model embeddings combined with point cloud annotation and clustering. We show that not only is IF-SitePred competitive with state-of-the-art methods when predicting binding sites on experimental structures, but it performs better on proxies for novel proteins where low accuracy has been simulated by molecular dynamics. Finally, IF-SitePred outperforms other methods if ensembles of predicted protein structures are generated.

## Introduction

A key part of early-stage drug development is building a thorough understanding of the protein target of interest. Identification of potential ligand-binding sites facilitates a host of techniques, such as hit identification, small molecule screening, functional prediction, off-target binding prediction and binding site comparison [1, 2]. Many methods have been developed to locate ligand-binding pockets on protein structures. Originally, these methods were designed for use on experimentally determined structures, but the development of AlphaFold [3] and other accurate protein structure prediction tools [4, 5, 6] now allows the exploration of the three-dimensional features of the protein, including predicting or identifying the ligand binding site from structural models.

### Strategies for protein binding site prediction

Various strategies exist for protein binding site prediction. Where the protein of interest has close homologues that have already been crystallised with ligands, the protein binding site can be inferred based on the alignment to these complexes [7, 8, 9]. However, if no homologous structural data is available, predictions must be made in other ways. Sequence-based methods use the amino acid sequence of the protein which is sometimes enriched with predicted structural features such as secondary structure, solvent accessibility or hydrophobicity. However, these methods achieve lower success compared with structurally-informed methods, and so they tend to be used for the prediction of specific ligand binding sites, such as carbohydrates [10]. Structurally-informed methods include geometry-based [11] and probe-based [12] methods, which use the shape of the protein surface to predict which regions are likely to bind ligands. While many new machine learning-based methods have been developed recently, there are some non-machine learning-based methods that are still commonly used, such as FPocket [11], FTSite [12], and DoGSite3 [13]. FPocket uses Voronoi tessellations to facilitate alpha-sphere labelling and clustering, followed by partial least squares fitting for ranking of binding sites. FTSite places 16 small molecular probes over the surface of the protein and finds clusters of favourable positions using empirical free energy functions. DoGSite3 uses a difference of Gaussians algorithm for finding binding sites of specific ligands.

### Machine learning in protein binding site prediction

The move to machine learning-based methods is driven by the requirement to learn underlying patterns in large sets of data that have proved difficult to learn or represent using physics-based or geometry-based approximations [14]. Where a protein of interest has no ligand-bound homologues, binding site prediction requires an understanding of the specific chemical environment needed to bind a ligand. The complexities of protein structure, the large number of data points and the number of features that make up this environment make this an ideal problem to approach via machine learning functions. Additionally, making predictions using machine learning can be significantly faster than other techniques, allowing predictions to be made on large sets of proteins with a lower computational cost.

Proteins can be represented in a variety of ways to facilitate machine learning approaches, and can be split broadly into two groups: featurised representations and learnt representations [15, 16].

Featurised representations consist of extracted spatiochemical information such as atom type, residue type or euclidean coordinates, usually combined with calculated features such as solvent accessibility, secondary structure, hydrophobicity and charge [17, 18, 19]. These features are then represented by annotation of the atom or residue, usually as part of a point cloud, graph or voxelisation [20]. Alternatively, the protein surface itself can be represented using a mesh, Voronoi tessellation or pseudo-atoms annotated with features that describe their chemical environment. The chosen representation is then used to train a machine learning model that predicts on the same type of data. Examples of featurised methods include P2Rank [21], DeepPocket [22], BiteNet [23], DeepSurf [24], NodeCoder [25] and PUResNet [26].

Learnt, or non-featurised, representations commonly use a series of vectors to describe a protein. These are often weights from the final layer of a transformer architecture that has been trained to predict masked residues of a protein [27], and are often referred to as embeddings. They represent protein residues as continuous vectors rather than discrete variables, and describe the environment in which a residue exists with respect to neighbouring residues. By using these embeddings along with experimentally-determined labels, machine learning models have been trained to predict features [27] such as ligand binding [28], protein-protein interactions, disease variants [29] or structural features [16]. Unsupervised learning can also be conducted, and has been used for enzyme function prediction [30]. The previously mentioned embeddings generated by language models have previously been adopted to train a secondary model to identify ligand-binding residues [31].

However, these techniques did not incorporate three-dimensional structural information. In 2020, the geometric vector perceptron (GVP) architecture, which did contain this 3D information, was introduced to leverage the proteins’ geometric and relational aspects in a sequence recovery task [32]. This particular architecture was later combined with a generic transformer to create ESM-IF1 [33] which produces embeddings that consist of 512-dimensional vectors for each residue of a protein structure. The ESM-IF1 embeddings have been used for epitope prediction [34] and protein-protein interaction (PPI) prediction [35]. Successes such as these suggest that the embeddings contain task-relevant information relating to the protein function and biochemical activity. The embeddings do not take into account side chain positions, but only the protein backbone, which should make the embedding robust to errors in the side chain positioning. This should offer advantages when making predictions on low-accuracy structures, whether experimental or predicted.

### Binding site prediction on predicted structures

Most binding site prediction methods are designed to predict binding sites from the ligand-bound (holo) structure of the protein, however this is not necessarily the most useful task. Since mid-2021, highly accurate structure predictions for many proteins have been available, first from AlphaFold [3], and later from others, including RosettaFold [5], OmegaFold [6] and ESMFold [4]. This means that for many novel proteins, there is now a structural starting point for protein binding site prediction where previously there was no experimental information available.

To aid this process, AlphaFill [36] was developed following the release of AlphaFoldDB [37], a database of predicted protein structures. AlphaFill ‘fills in’ predicted protein structures with putative ligands by searching for areas of local sequence similarity between predicted proteins and existing complexes in the PDB [38]. An alignment and ‘transplantation’ strategy places ligands in potential binding sites of AlphaFold-predicted protein structures (from here referred to as AF2 structures). For novel proteins that have regions of at least 85 residues with higher than 25% sequence identity to proteins with ligands bound in the PDB, AlphaFill provides a first step to locating ligand-binding sites. Where sequence identity for a minimum of 85 residues is higher than 40%, binding site RMSD is rarely higher than 2Å, suggesting a good match. AlphaFill was able to fill over 59% of proteins in AlphaFoldDB in February 2022 with ligands. This leaves a large proportion of proteins that do not have close homologues already experimentally solved with ligands bound. These proteins will therefore require template-free binding site prediction.

### Binding sites of predicted protein structures

The most commonly used accuracy measurement for protein structure predictions is the global RMSD of their backbone atoms when aligned to the experimentally-solved structure (from here referred to as the PDB [38] structure); a value below 2Å is taken to indicate the model is of high quality. However, this global measure does not provide an assessment of the local quality of the binding site in this predicted structure.

Binding site prediction methods have already been applied to AF2 structures. FPocket [11] was used to compare volumes of binding pockets in PDB structures and their high-accuracy AF2 counterparts (median all-atom RMSD of 1.54Å), and a 20% reduction in binding pocket volume in AF2 structures was found [39]. A 2022 study on AF2 structures showed that binding site prediction by AutoSite [40] was much less successful where mean residue confidence (pLDDT) for a protein was below 90%, with F-scores reduced by around 80% compared with predictions on holo, apo or high-confidence AF2 structures [41]. The same study also found that only 25% of residues are predicted with confidence over 90%, indicating that many AF2 structures may be difficult targets for binding site prediction.

Several studies on docking small molecule ligands into AF2 structures have been published [39, 42, 43]. Despite high accuracy in the test structures (17 of 22 had RMSD lower than 2Å in [43]), docking proved much more difficult for AF2 structures than their PDB counterparts [43]. Even when controlling for the accuracy of the predicted structure around the binding site, docking remained a challenge: a recent study found that even proteins with binding site all-atom RMSD as low as 1Å were significantly more difficult to dock into than experimentally-determined structures [44].

These studies all suggest that even accurate predictions may exhibit significant differences in binding sites. However, in a recent paper [21], P2Rank was found to have similar levels of accuracy in predicting binding sites on several thousand AF2 structures and PDB structures of the same proteins on two different test sets (HOLO4K and COACH420).

### Accuracy of predicted protein structures

The proteins in commonly-used test sets for binding site prediction (such as COACH420 and HOLO4K) are by definition publicly available as protein-ligand complexes. Therefore, it is possible they contain many proteins that are in the training sets for protein structure predictors [41]. This would result in predicted structures being much closer to the PDB structure than would be achieved for novel targets without existing close homologues. The consequence of this would be that the datasets used to test tools on ‘predicted structures’ would not be representative of predicted structures of novel targets, limiting the effectiveness of any evaluation.

The Critical Assessment of Structure Prediction (CASP) [45] carries out rigorous blind testing of protein structure prediction methods and evaluation of results by independent assessors [46]. For the CASP iteration in which AF2 was first present (2020), the best predicted structure for each target was taken and the fractions of targets predicted at different levels of accuracy were calculated. This provides insight into the accuracy of current protein structure prediction methods based on the availability of structural information of related proteins. Proteins are grouped based on the availability of related proteins with existing structures: of the ‘Free Modelling’ (FM) group that have no detectable homology to existing protein structures, over 50% of proteins present in the CASP test set have RMSD values greater than 2Å when aligned to the experimentally-solved structures. These are the proteins that cannot be filled by AlphaFill, so will require template-free binding site prediction.

One of the recent studies looking at docking into AF2 structures specifically selected proteins that were not in the AF2 training set for their test set [44], and found that most of these structures had between 2 and 4Å all-atom RMSD when compared to the experimental structures, further confirming that novel proteins which are targets for template-free binding site prediction are expected to have RMSD values in this range. Therefore, we would expect template-free binding site prediction tools to be evaluated using AF2 structures that have up to 4Å all-atom RMSD.

### Protein dynamics and binding sites

Protein structures, whether experimentally-determined or predicted, represent just a single snapshot of dynamic systems. This can limit prediction success, as the protein pocket may not be present in the particular structure that is being used for prediction. An analysis of BiteNet’s predictions on a minimization molecular dynamics (MD) trajectory of an adenosine A2A receptor showed that an allosteric site became detectable by BiteNet at a backbone global RMSD value of just 0.4Å compared to the original structure [23]. This emphasises how significant changes can happen to the structure of the binding sites at a local scale even when the change is negligible on a global scale. While this has importance in prediction of binding sites of experimental structures, it is even more relevant when using predicted protein structures, as ‘highly-accurate’ structures (*<*2Å to the experimental structure) may contain large local differences that make it difficult to identify any binding sites correctly.

Currently, MD simulations remain the only proven way of generating multiple structures for protein binding site prediction but their usefulness is limited because these simulations are computationally costly. Computationally cheaper generation of multiple protein conformations is an area of much interest [47, 48, 49, 50, 51], however, it is not yet clear if these methods are able to replace effectively information gained from MD simulations.

### Summary

Here we describe IF-SitePred, a method for protein binding site prediction using representations obtained from ESM-IF1 [33]. We compare the performance of our method to other commonly-used binding site prediction tools on PDB structures taken from the HOLO4K test set and their equivalent AF2 structures. We assess the accuracy of the predicted structures, and use molecular dynamics simulations of predicted structures to create lower accuracy protein models (up to 4Å RMSD) and evaluate how binding site prediction success varies with structural accuracy. Finally, we show that by using ensembles of structure predictions, the prediction of binding residues can be greatly improved. We find that IF-SitePred achieves superior binding site prediction compared to commonly-used methods on low-accuracy protein structures, particularly where multiple structures are available.

## Methods

### Datasets

We selected HOLO4K as our test set to facilitate comparison to other methods. For 4309 proteins in the HOLO4K set, we used the UniProt [52] ID mapping service to map each PDB code to the AlphaFold Protein Structure Database [37]. For the 3914 proteins that appeared to have a corresponding AF2 structure, we verified that the sequence identity between the PDB structure and the prediction was over 90% (100% was not always possible due to the presence of tags or absence of flexible regions in the PDB structure). This removed 1636 protein pairs, leaving 2278 proteins with correctly matched sequences. We then clustered each pair of sequences using MMseqs2 [53] to ensure our test set did not contain any pairs of proteins more similar than 90%. This resulted in 691 viable pairs of PDB and AF2 structures. To make it possible to evaluate predictions on the AF2 structures, we aligned each prediction to its corresponding PDB structure. Just 14 of the 691 predictions had a backbone RMSD above 2Å and each of these were visually inspected to check whether the binding sites aligned well enough to be included in the analysis. Of these, 11 were retained, including pairs with RMSD values up to 16Å (these contained some large differences in relative domain positions compared with their PDB counterparts, but still had well-aligned binding sites). We named the final set of 688 pairs the HOLO4K-AlphaFold2 Paired (HAP) set. We also extracted a set of 280 pairs which only contained proteins with lower than 25% sequence identity (calculated using Diamond [54]) to the P2Rank [17] training set (referred to as the HAP-small set).

Our training set consists of structures taken from Binding MOAD [55, 56, 57]. The Binding MOAD platform contains 11058 families (clusters) of proteins with each cluster containing a leader (the cluster centre) and members which each have over 90% sequence identity to the leader. We first removed any family for which the leader had greater than 25% sequence identity to any protein in our test set. For each of the remaining 6550 families, we aligned all members to the family leader, and labelled the residues of the leader as follows: any residues within 5Å of the ligand of the leader were labelled as binding residues; any remaining residue within 5Å of any ligand bound to member structures were not used in the dataset, to avoid the potential for false negative annotations; the remainder were labelled as non-binding. Only residues with relative surface accessibility over 0.02 (calculated using the PyMOL API [58]) were included. This resulted in 143,022 binding residues and 1,414,153 non-binding residues for use in training.

### Model training

For each protein in the training set an ESM-IF1 embedding [33] was generated. The residue annotations were applied as above, and the residues were treated independently. Each training set was balanced, made up of a random 80% sample of the binding residues and an equal number of randomly sampled non-binding residues. Using a bootstrapping sampling method with replacement, we generated 40 training sets. We initially used AutoML [59] to train models on all 40 datasets and found that in the majority of models, the LightGBM model [60] had the highest performance on the validation data (randomly taken from the training set). We therefore trained LightGBM models for all datasets, using a 10% random sample of the input data as a validation set. For all models, the parameters were fixed. A binary objective was used, along with a ‘binary_logloss’ metric parameter. Based on commonly selected parameters in the AutoML models, gradient-boosted decision trees (GDBT) were used, with no feature pre-filtering and no early stopping round; the number of leaves was set at 200 and 200 iterations were used.

### Binding site prediction with IF-SitePred

For a protein in the test set, an ESM-IF1 embedding was generated and each residue was independently predicted by each of the 40 models to be ligand-binding or non-ligand-binding, with a minimum predicted probability of 0.5 for positive labelling. Only if all 40 models predicted a residue to be binding was a positive label applied. Using the PyMOL API [58], a point cloud on a 1.5Å grid was generated around the protein, containing only points between 3 and 6Å from any protein atom. For every residue that was labelled as binding, points within 4.5Å of the residue were saved. Points that were saved three or more times (i.e. were within 4.5Å of three or more residues labelled as binding) were clustered using the DBSCAN algorithm from Scikit-learn [61], using a 1.7Å cutoff to separate clusters. Clusters were ranked simplistically, using the total number of points in the cluster (including repetitions of the same point), and the centres of the top-ranked clusters were calculated by taking the mean coordinates of all points. This process is summarised in Figure 1. All distance thresholds in the point cloud labelling and clustering were selected to maximise success on training set proteins where all residues were labelled correctly (99% success when predicting top 1 pocket). To explore how small changes in these thresholds impact prediction success, we adjusted the thresholds higher and lower within 1Å for all values except for the clustering distance threshold, which was adjusted by 0.1Å (Table S1).

**Figure 1:**
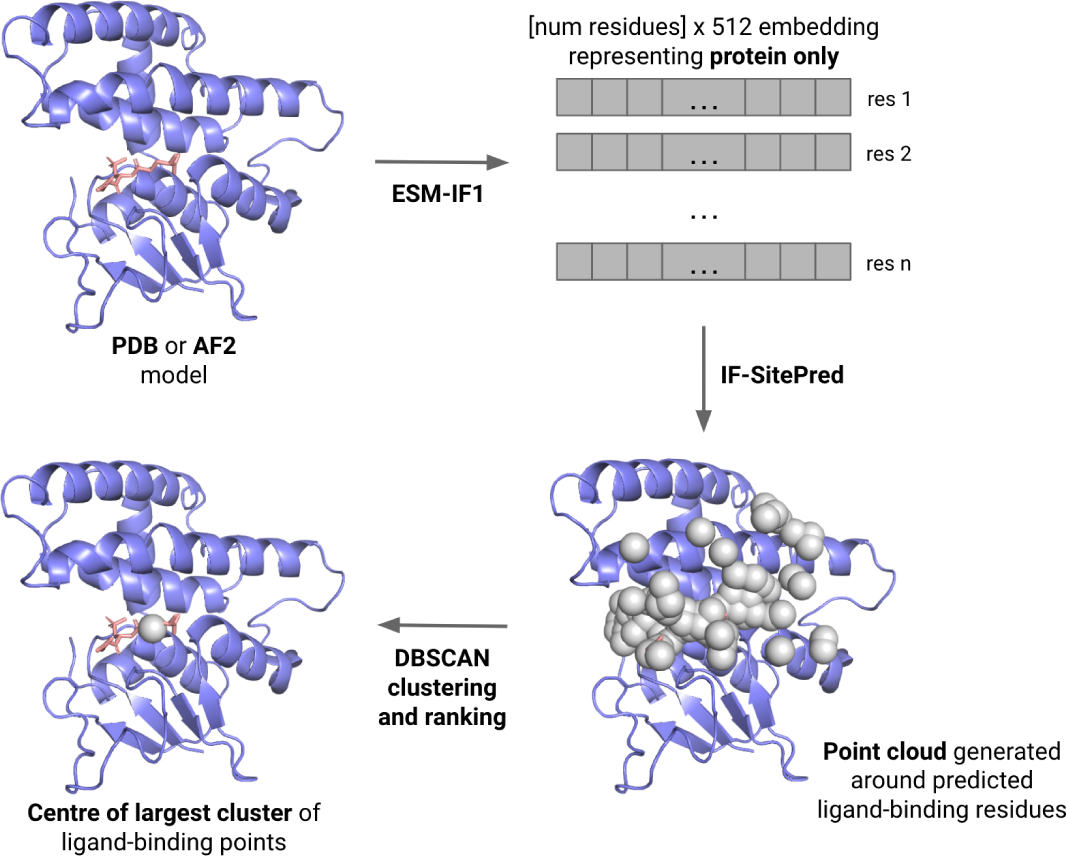
Summary of IF-SitePred and binding site centre calculation. ESM-IF1 embeddings were generated for each residue of the protein, and the model labelled each of these as binding or non-binding. A point cloud around predicted binding residues was generated, made up of points that were within 4.5Å of at least three predicted binding residues. These points were clustered using the DBSCAN algorithm. Clusters were ranked in order of number of points, and the mean coordinates of each cluster was taken as the binding site centre.

When predicting on an AF2 structure, the predicted structure was first aligned to its PDB counterpart prior to removal of the relevant chain of the PDB structure. The point cloud generation and clustering protocol was then applied as above.

For comparison of predictions with a random baseline, the number of residues predicted as ligand-binding was calculated, and the same number of surface residues were randomly annotated as binding, keeping the same proportions of sub-surface (relative surface area between 0.01 and 0.05) and surface (relative surface area over 0.05) residues. An identical protocol to that above was used for point cloud generation and clustering.

### Evaluation Criteria

Several evaluation strategies have been used for binding site prediction. The traditional metrics of area under the receiver operator characteristic curve (AUROC) or accuracy are not appropriate for imbalanced problems as very high scores can be achieved by predicting all residues as non-binding [62]. DCA, DCC, DVO or atom IoU (Table 1) are commonly used, however DCC, DVO and IoU are based on the assumption that the ligand in the PDB complex is the perfect ligand. While this may be the case, it is not certain, and so we opted to use DCA to evaluate and compare our prediction method, where we measure success by whether the centre of the predicted site is within 4Å of any ligand heavy atom. This avoids reliance upon an over-specific definition of the binding site. Several methods calculate DCA for the top-ranked pocket and top-n ranked pockets (where n is the number of ligands bound to the target protein), however this assumes that the experimental complex contains all possible correct ligands, and so we opted to use the top-ranked pocket and the top-3 ranked pockets.

**Table 1:** Commonly-used evaluation metrics for protein binding site prediction.

| Metric | Definition |
| --- | --- |
| DCA | Distance from site Centre to ligand Atom: the distance in Å from the predicted binding site centre to any ligand heavy atom from the experimentally-determined complex |
| DCC | Distance from site Centre to ligand Centre: the distance in Å from the predicted binding site centre to the centre of the ligand from the experimentally-determined complex |
| DVO | Discretized Volume Overlap: intersection over union of the volume of the predicted binding site and the volume of the ligand from the experimentally-determined complex |
| atom IoU | atom Intersection over Union: for use in methods where protein atoms are individually labelled as ligand-binding or non-ligand-binding, intersection over union of the predicted ligand-binding atoms and the experimentally-observed ligand-binding atoms is calculated |

As a secondary evaluation method, we used F1 score of residue annotation as ligand-binding or non-ligand-binding. F1 score takes into account precision and recall, thus avoiding the issue created by the imbalanced number of positive and negative labels in the data. A score of 1 indicates perfect prediction, however predicting all residues as non-binding would yield an F1 score of 0.

### Comparison to existing methods

We selected three popular methods for comparison, based on the range of techniques used, the availability of command-line software and the compatibility of their training sets with our test set: FPocket [11], P2Rank [17] and DeepPocket [22]. FPocket uses Voronoi tessellations and alpha-sphere clustering for prediction to find clefts on the protein surface that have the correct size and shape for ligand binding. P2Rank annotates points on the solvent-accessible surface area of the protein based on feature-vectors applied to exposed protein atoms, labels each point as ligand-binding or non-ligand-binding using a random forest classifier, and ranks sites according to their cumulative ligand-binding score. DeepPocket is a 3D-CNN based method which re-scores pocket centres identified by FPocket, then elucidates the shapes of the predicted pockets.

For all four methods, we compared binding site prediction success as defined by DCA on the top-ranked pockets and top-3 ranked pockets for PDB and AF2 structures for both the HAP and HAP-small sets, as defined above.

For FPocket and DeepPocket, default parameters were used for binding site prediction on all structures. For FPocket, the centre of each binding site was calculated by taking the mean coordinates of all points in the pocket. For P2Rank, default parameters were used for PDB structures, and for predicting on AF2 structures, the AlphaFold-specific configuration was used.

### Ligand similarity to training sets

Similar to the dataset splitting strategy for preventing protein sequence bias, we also checked whether the methods displayed a bias towards the ligands present in their training sets as follows. Ligand similarity was calculated using RDKit [63] USRCAT [64] similarity. For each test protein, the USRCAT similarity for each ligand bound was calculated for every ligand bound to a training set protein. The highest value was taken and the top-1 success rate for proteins at varying levels of ligand similarity was compared with the mean top-1 success rate. This analysis was performed for each of IF-SitePred, FPocket, P2Rank and DeepPocket with their respective training sets.

### Molecular dynamics simulations for testing predictions on low-quality structures

The AF2 structures in the HAP set are highly accurate, with only 11 of 688 predictions having backbone RMSD values over 2Å to the PDB structure. This represents just 1.6% of the predictions, whereas the analysis of the best prediction for each target in CASP14 [46] suggested that for free modelling targets (those targets without a known structural homologue), over 50% of predictions are likely to have an RMSD over 2Å. Additionally, a study on GPCRs found that most structural predictions that were not in the AF2 training set had RMSD values between 2 and 4Å [44]. To explore how binding site prediction success changes as predicted structures become less accurate, we filtered the HAP set for physiologically monomeric proteins with fewer than 230 amino acids, resulting in a set of 21 proteins for which we conducted MD simulations to generate structures that were representative of low-accuracy protein structure predictions.

We used the AF2 structures of each protein for the MD simulations. For each AF2 structure, pKa values were estimated using h++ [65, 66, 67] to assign residue protonation states. Ions were added using tleap [68] to neutralise the system. We then used the Amber [68] protein forcefield (fF14sb) within OpenMM [69] to heat the proteins from 298K to 548K, with the system simulated for 20ns for each 10K interval. Using MDAnalysis [70], the trajectory was randomly sampled to extract 10 structures for each 0.25Å interval from RMSD values up to 8Å when aligned to the PDB structure.

### Evaluation

For IF-SitePred, P2Rank and DeepPocket, we predicted binding sites for structures with RMSD values up to 7Å from the PDB structure. Mean rates of top-1 success and top-3 success for each 0.25Å interval were calculated for each method.

### Combining predictions by using multiple models on multiple structures

We tested two ensembling methods. For the first, we trained 40 models on different samples of the training data, and combined the results of these to make our final predictions. For the second, we made predictions on multiple protein structures (these could be generated by different protein structure prediction tools or by molecular dynamics) and combined these.

We tested the improvements made by using multiple models and multiple medium-accuracy protein structure predictions. The protein structures we used to test this were from MD simulations of AF2 structures with RMSD to the PDB structures lower than 4Å, as this would cover around 82% of predictions on free-modelled proteins in CASP14 [46].

We implemented four prediction pathways to understand the effects of using single or multiple models on single or multiple structures (Figure S1). Residues were annotated as ligand-binding or non-ligand-binding by either 40 models (as previously) or by just one model, and point clouds were generated and clustered as previously. The top 3 binding sites for each structure were used to re-annotate only the residues within 5Å of their points as ligand-binding, as this is a commonly used distance threshold for intermolecular interactions. To view the effect of combining predictions on multiple structures, 9 other MD-generated structures of the same protein were randomly selected, and only residues that were predicted as ligand-binding in at least 3 of the 10 structures were given a final prediction as ligand-binding. This minimum threshold of 3 positive predictions for a residue was arbitrarily selected. The F1 scores for these final predictions were calculated to compare residue annotation for these four prediction pathways.

Additionally, we combined residue labels of 10 structures (as above) 1000 times for 150 randomly-selected frames of each of the 21 proteins for which MD had been performed using IF-SitePred, P2Rank and DeepPocket to compare which of these methods are able to benefit from this ensemble method.

## Results

### IF-SitePred binding site prediction is competitive with state-of-the-art methods

The prediction of ligand-binding sites on the surfaces of proteins is a useful step towards understanding the function and druggability of novel targets. It is particularly important in the era of accurate protein structure prediction, where we often have a predicted structure before any experimental studies have been carried out. We developed IF-SitePred, a binding site prediction tool which avoids featurisation, and instead uses embeddings generated by ESM-IF1 as the basis for labelling protein residues as ligand-binding or non-ligand-binding. This is followed by point cloud annotation and clustering to determine the three most likely binding sites and their centres. To evaluate our method, we predicted binding sites on hundreds of proteins in the HAP set and compared our rate of success to that of FPocket, P2Rank and DeepPocket. In particular, we compared prediction success on experimental (PDB) structures with those predicted by AlphaFold (AF2). To ensure that we evaluated the tools on a range of proteins that were sufficiently different to the training data from each method, we designed the HAP set to include 688 proteins (both PDB and AF2 structures) which have no more than 90% sequence identity to any other protein in the set and no more than 25% sequence identity to the training sets of IF-SitePred, FPocket and DeepPocket. The HAP-small set is a subset of 280 proteins from the HAP set, made up of proteins that have no more than 25% sequence identity to the training data used for comparator method P2Rank. Prediction success was measured by top-1 and top-3 DCA, which checks whether the predicted binding site centre is within 4Å of any heavy atom of the experimentally-determined ligand. This measure avoids the assumption that the observed ligand perfectly fills the site.

The binding site prediction success rates for IF-SitePred, FPocket, P2Rank, and DeepPocket are shown in Table 2. On PDB structures, all methods performed similarly well. On the HAP set, DeepPocket achieved the highest top-1 success rates, but was outperformed by IF-SitePred on top-2 and top-3 success rates. FPocket had the lowest success rates. By using the HAP-small set to compare methods, we observed that P2Rank outperformed all other methods at top-1 success, but shared top performance with IF-SitePred when considering top-3 success. Overall, P2Rank had similar performance on the HAP and HAP-small sets, suggesting the method generalises well. Similar results were observed for IF-SitePred, DeepPocket and P2Rank when predicting binding sites on AF2 structures, with these three methods sharing the highest success rates across the HAP and HAP-small sets. However, FPocket experienced a significant drop-off in performance on AF2 structures. Given that DeepPocket is a prioritisation method that takes FPocket’s predictions as an input, this suggests that the ranking procedure used by FPocket failed on AF2 structures, as opposed to FPocket having difficulties in identifying the ligand-binding sites on the protein’s surface. This could be due to the lower pocket volume of AF2 structures [39].

**Table 2:** Success rate of binding site prediction of IF-SitePred and commonly used existing methods. IF-SitePred, P2Rank and DeepPocket are competive across PDB and AF2 structures, whereas FPocket experiences a significant loss of performance on AF2 structures. Success rates of top-1, top-2 and top-3 binding site prediction as measured using DCA is shown, where success is defined as the centre of the predicted binding site being within 4Å of any ligand heavy atom. We show results for IF-SitePred, FPocket, P2Rank and DeepPocket on two test sets that contain PDB and AF2 structures respectively. For each test set, the highest success rate is shown in bold, and the lowest success rate is shown in italics.

| HAP: 688 proteins |  |  |  |  |  |  |  |  |  |  |  |
| --- | --- | --- | --- | --- | --- | --- | --- | --- | --- | --- | --- |
| PDB | baseline | IF-SitePred | FPocket | P2Rank | DeepPocket | AF2 | baseline | IF-SitePred | FPocket | P2Rank | DeepPocket |
| Top 1 | 0.12 | 0.76 | *0.75* | **0.81** | 0.78 | Top 1 | 0.09 | 0.77 | *0.50* | **0.81** | 0.78 |
| Top 2 | 0.17 | 0.89 | *0.81* | **0.90** | 0.87 | Top 2 | 0.17 | **0.89** | *0.60* | 0.88 | 0.87 |
| Top 3 | 0.22 | **0.93** | *0.83* | **0.93** | 0.89 | Top 3 | 0.20 | **0.94** | *0.67* | 0.89 | 0.90 |

Table 2: IF-SitePred, P2Rank and DeepPocket are competitive across PDB and AF2 structures, whereas FPocket experiences a significant loss of performance on AF2 structures.
| HAP-small: 280 proteins |  |  |  |  |  |  |  |  |  |  |  |
| --- | --- | --- | --- | --- | --- | --- | --- | --- | --- | --- | --- |
| PDB | baseline | IF-SitePred | FPocket | P2Rank | DeepPocket | AF2 | baseline | IF-SitePred | FPocket | P2Rank | DeepPocket |
| Top 1 | 0.12 | 0.75 | *0.73* | **0.78** | 0.75 | Top 1 | 0.10 | **0.76** | *0.48* | **0.76** | 0.75 |
| Top 2 | 0.18 | **0.89** | *0.80* | 0.86 | 0.85 | Top 2 | 0.17 | **0.88** | *0.58* | 0.85 | 0.86 |
| Top 3 | 0.24 | **0.92** | *0.82* | 0.91 | 0.88 | Top 3 | 0.20 | **0.94** | *0.65* | 0.88 | 0.90 |

For proteins where IF-SitePred failed to make a successful top-1 prediction on the PDB structure, around half were also failures when using the AF2 structure. However, the other half were successfully predicted when using the AF2 structure, suggesting that predicted structures sometimes contain information about ligand binding that is not present in the ligand-bound PDB structure. A similar result was found in the recent P2Rank paper [21].

### Accurate binding site prediction is not dependent on ligand similarity to the training set

By removing any protein from the training set with more than 25% sequence identity to any test set protein, we attempted to ensure that our predictions were not based on the model learning the sequences from the training set. However, the model could be learning ligand-based information that was contained in both the training and test sets. To explore this possibility, we investigated whether we were able to predict binding sites more successfully on proteins that have similar ligands to our training set.

For each protein-ligand complex in the HAP-small set, we calculated the maximum ligand similarity to any ligand that bound to proteins in the training set for each of IF-SitePred, FPocket, P2Rank and DeepPocket. Ligand similarity was defined using USRCAT fingerprints, which takes into account the ligand shape and pharmacophoric features. These values were then compared to the mean top-1 success (found in Table 1). These results are shown in Table 3.

**Table 3:**
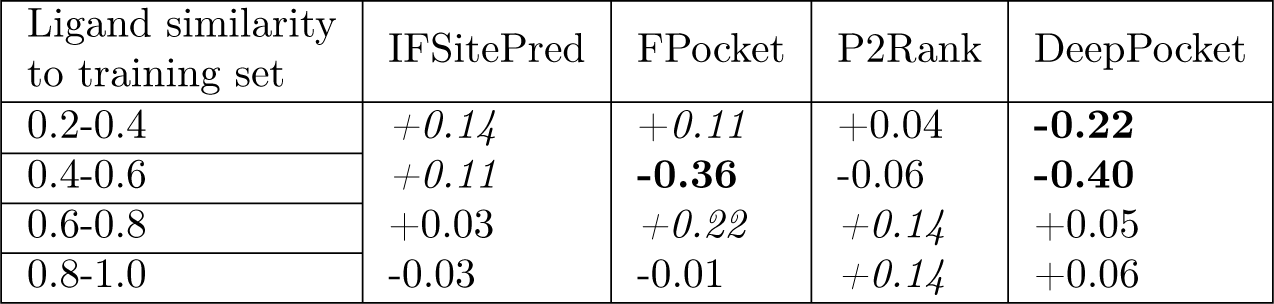
DeepPocket performs worse than average on sites that bind ligands significantly different to the training set. We calculated the USRCAT similarity for the most similar ligand in the training set to that binding each protein in the HAP-small set, and calculated the fraction difference between overall top-1 success rate (from Table 1) for each 0.2 interval of USRCAT similarity. Success rates differing by over 10% from the mean value are shown in bold (performance loss) or italic (performance gain).

| Ligand similarity to training set | IFSitePred | FPocket | P2Rank | DeepPocket |
| --- | --- | --- | --- | --- |
| 0.2-0.4 | *+0.14* | *+0.11* | +0.04 | **-0.22** |
| 0.4-0.6 | *+0.11* | **-0.36** | -0.06 | **-0.40** |
| 0.6-0.8 | +0.03 | *+0.22* | *+0.14* | +0.05 |
| 0.8-1.0 | -0.03 | -0.01 | *+0.14* | +0.06 |

If our predictions were dependent on ligand similarity, we would observe that where ligands bound to the binding site are significantly different to the training set, the binding sites would be predicted with lower success than those where similar ligands were present in the training set. We did not see this trend in the IF-SitePred results, so we can be confident that our predictions do not rely on ligand similarity to the training set. P2Rank also does not exhibit a significant bias towards proteins that have ligands that bound to training set proteins. However, where a site binds a ligand with only dissimilar ligands in the training set (USRCAT similarity lower than 0.4), DeepPocket and FPocket are significantly less able to correctly predict the binding site at the highest rank.

In our tests IF-SitePred and P2Rank are not affected by similar ligands being available in the training set, whereas DeepPocket and FPocket are, meaning that they may not generalise well to the prediction of binding sites of novel ligands.

### IF-SitePred outperforms other methods in top-3 binding site prediction on MD structures

When verifying the binding site alignment of the AF2 counterparts of the test set proteins, we observed that only 11 proteins in the final HAP set had backbone RMSD to the PDB structure above 2Å (Figure 2). This represents just 1.6% of the predictions, whereas analysis of CASP14 results [46] suggested that for free modelling targets, over 50% of predictions had an RMSD over 2Å, over 30 times what we observe in our dataset.

**Figure 2:**
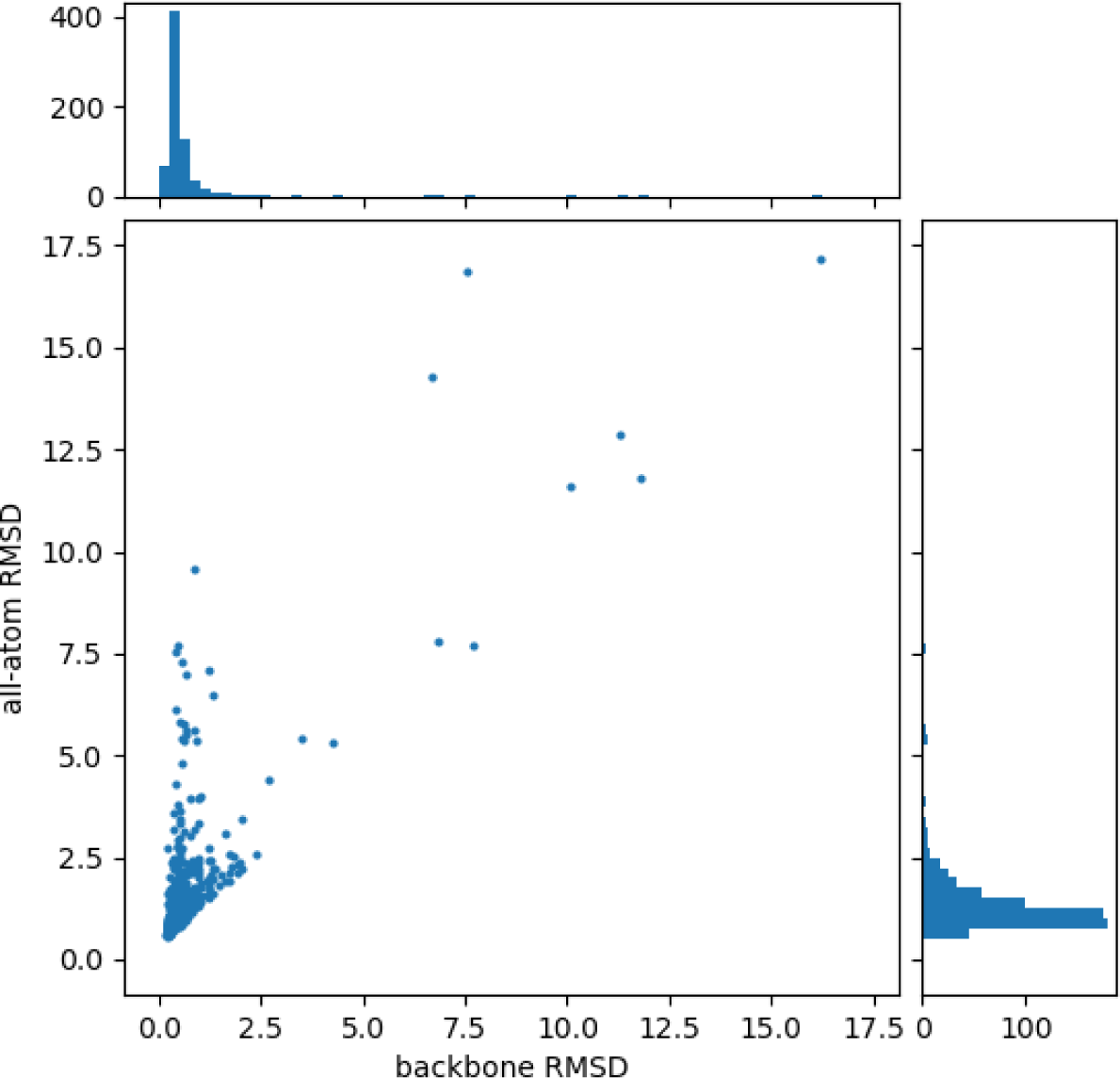
RMSD of AF2 structures in HAP set. AF2-predicted structures in the HAP set are extremely accurate compared with most free modelling AF2 predictions on novel proteins. Backbone RMSD is shown on the x-axis, and all-atom RMSD is shown on the y-axis. Each axis has a corresponding histogram to show the spread of values. Over 98% of AF2 structures in the HAP set have backbone RMSD values below 2Å.

To explore how binding site prediction success changes when the input structures are less accurate, we selected 21 monomeric proteins with fewer than 230 amino acids from the HAP set for which we conducted molecular dynamics simulations, followed by binding site prediction.

We heated each protein from 298K to 548K, simulating for 20ns at each 10K interval, and sampled structures from every tenth frame. Structures were sampled uniformly up to 8Å RMSD (when aligned to the PDB structure) and binding sites were predicted using IF-SitePred, P2Rank and DeepPocket. Using DCA, the probability of success for each 0.25Å RMSD interval was calculated for top-1 and top-3 ranked binding sites (Figure 3). The means of these probabilities for each interval were calculated, and a line of best fit was determined. Even at just 1Å RMSD, performance was significantly reduced for all methods, with a reduction in success rates of up to 15% compared to the PDB or original AF2 structures.

**Figure 3:**
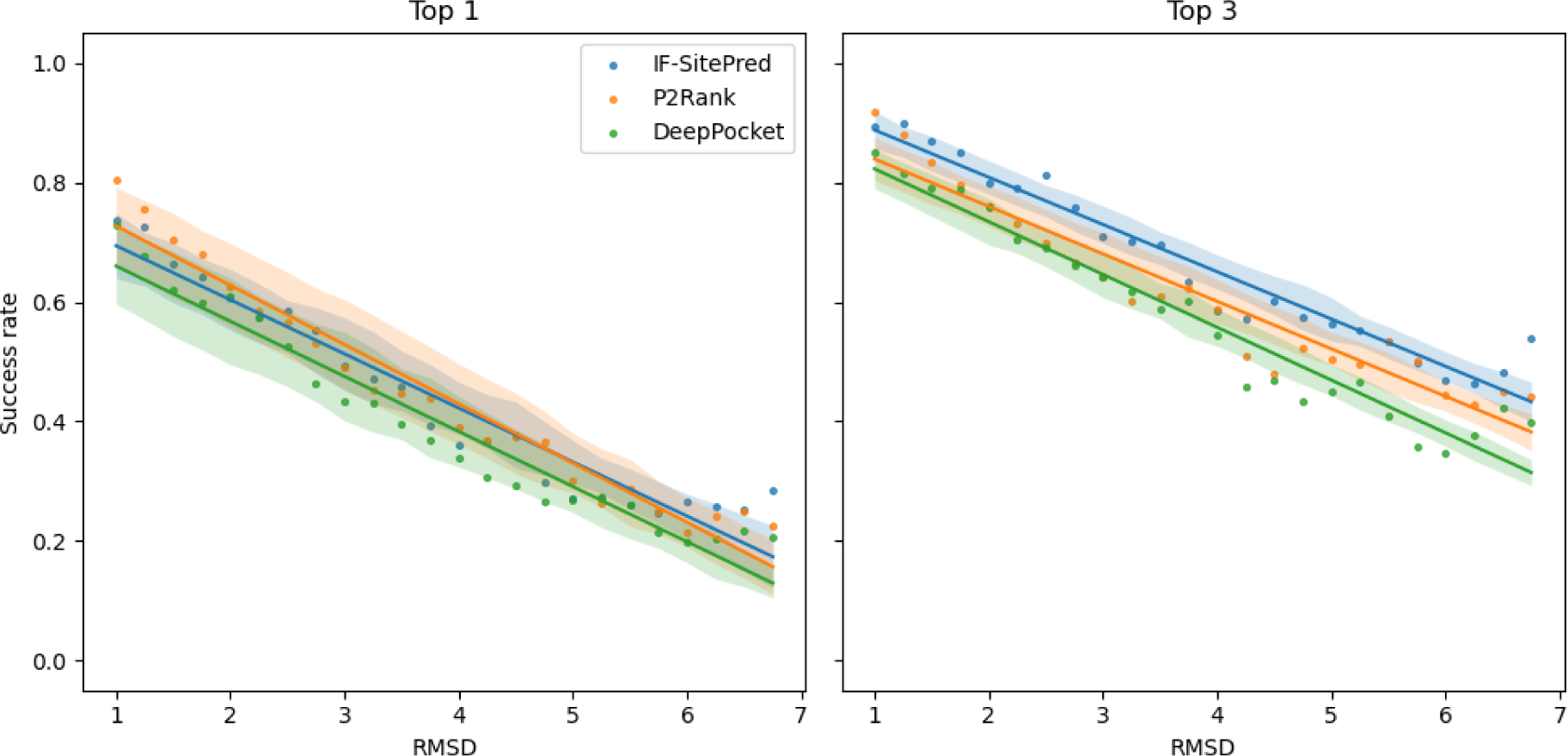
Binding site prediction on structures with decreasing accuracy. IF-SitePred outperforms P2Rank and DeepPocket when considering top-3 success on low-accuracy structures. We predicted binding sites on MD structures at increasing RMSD when aligned to PDB structures. Points represent mean success rate at each RMSD interval (value to value plus 0.25Å) across all 21 targets. Based on these, lines of best fit are calculated and plotted (solid lines of identical colour). Standard error of the mean across all targets is represented by the shaded areas. The three methods performed similarly when considering only the top-ranked binding site (left), with P2Rank performing slightly better at low RMSD values. However, IF-SitePred achieved higher top-3 success than P2Rank and DeepPocket at all RMSD values (right).

When comparing the three methods, we found that P2Rank performed best on structures up to 3Å RMSD, but had a greater loss of performance than IF-SitePred, which attained higher success rates at RMSD values over 4Å. When evaluating top-3 success rates, IF-SitePred achieved higher success rates than P2Rank and DeepPocket across almost all RMSD values. The difference grew at higher RMSD values, with IF-SitePred able to succeed 60% of the time on structures with an RMSD of 5Å, compared to 50% and 47% success for P2Rank and DeepPocket respectively. These results all suggest that IF-SitePred is more robust to errors in the protein structure.

### Combining predictions of multiple models on multiple structures improves predictive power

IF-SitePred uses 40 models that were trained independently on 40 different samples of the training data. Due to the imbalanced nature of the number of binding and non-binding residues, this allows the different models to learn the features of different sets of the non-binding residues, thus providing a final prediction informed by more data. Another strategy to maximise use of available data is possible when multiple structures of the same protein are available, whether by generating multiple structure predictions when using a tool like AlphaFold, or by performing molecular dynamics simulations: predictions can be made on multiple structures and combined to give a final prediction.

We implemented four pathways which used a combination of different models and structures (see Methods) to understand the improvements made when combining different predictions. For each prediction on an MD structure with RMSD values lower than 4Å, we calculated the F1 score of our predicted binding residues compared with the binding residues observed in the PDB protein-ligand complex (Figure 4).

**Figure 4:**
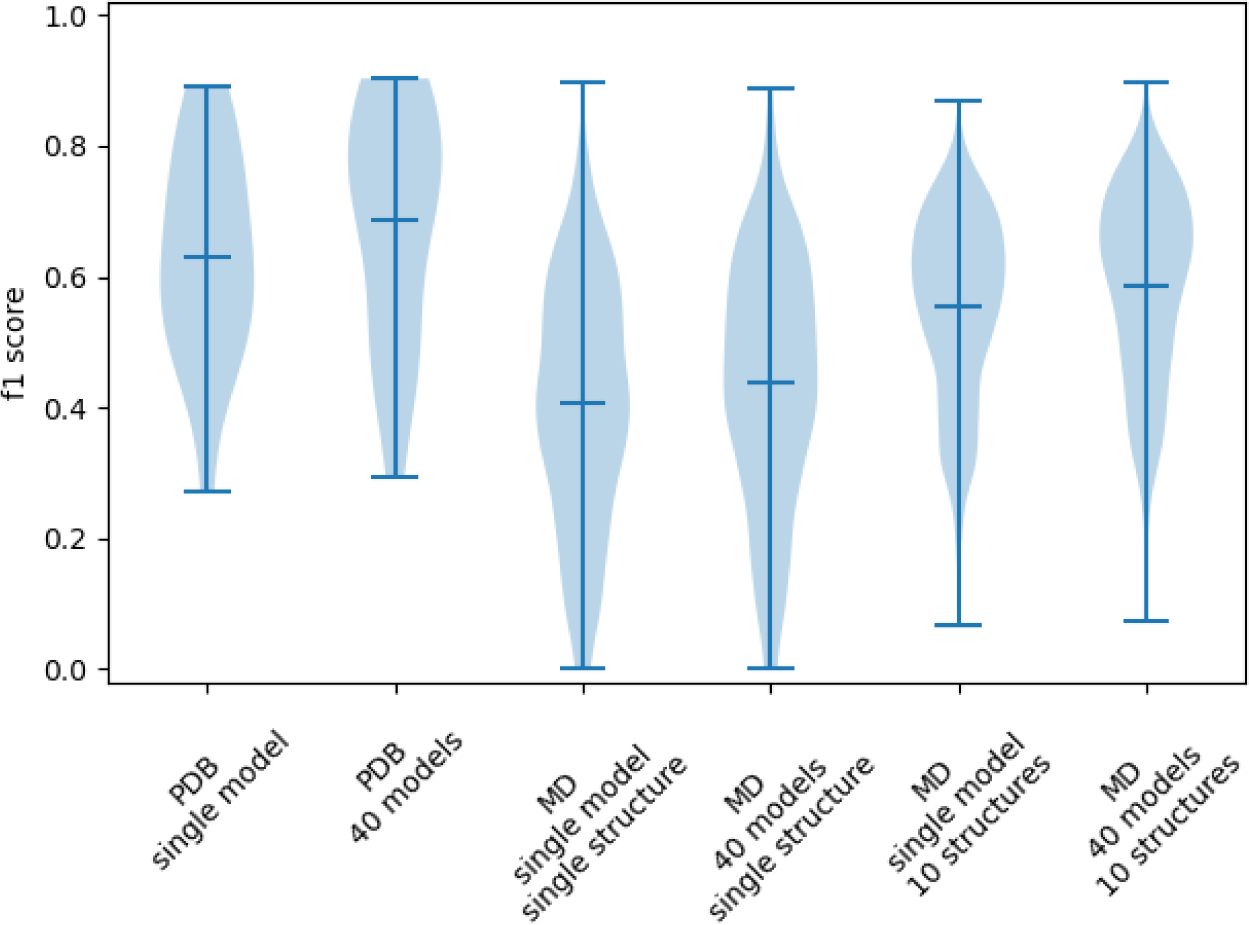
Combining predictions of multiple models on multiple structures: IF-SitePred. Using multiple predictive models on multiple protein structures improves IF-SitePred’s pre-dictive power. We compare F1 scores of predictions of binding residues for PDB and MD structures of 21 proteins when using single or multiple predictive models on single or multiple structures (MD only). Combining predictions of multiple models on multiple structures yielded the most accurate prediction of binding residues on MD structures, with F1 scores approaching those of predictions on PDB structures.

We first calculated the mean F1 scores of binding residue prediction on PDB structures using a single predictive model (0.62) and using 40 predictive models (0.68). When a single predictive model predicted the binding residues of single low-accuracy protein structures, the mean F1 score was just 0.41. When multiple models were used for the prediction, the mean F1 score rose to 0.43, and a further improvement to 0.59 was seen when multiple protein structures were also included. This represented an improvement of 44% when using two ensemble strategies compared to when only using single models and structures, and showed that by developing methods that take into account as much data as possible, we were able to mitigate errors in the data and make good predictions on flawed data that were significantly closer to the accuracy seen when predicting on high-quality data.

We additionally compared the benefits of using many structures to make final predictions for IF-SitePred, P2Rank and DeepPocket. This involved taking predictions from a single structure and comparing the F1 score with the combined prediction of the same structure with nine additional randomly-selected structures. This was repeated 1000 times for each target to ensure that the results were as representative as possible of each method. While the F1 scores across methods do not directly correspond to DCA success rate as the protocols for determination and ranking of protein binding site centres differ, we were able to compare the impact of combining multiple sets of predictions between methods. Additionally, we used the frequency of prediction of each residue as ligand-binding to create a multi-structure prediction probability, and used this to plot precision-recall curves for each method.

When just one structure was used to make a prediction, IF-SitePred slightly outperformed P2Rank and DeepPocket, in agreement with DCA-based results (Figure 3). When predictions for 10 structures were combined, all three methods had improved predictive power, showing the importance of understanding multiple states of the protein. IF-SitePred was able to benefit the most, with F1 score increasing by around 34%, compared with 22% and 15% improvements from P2Rank and DeepPocket respectively (Figure 5(a)). We expect that this would translate to a higher DCA success rate for IF-SitePred compared to P2Rank and DeepPocket when predicting binding sites on error-prone structures. Additionally, we show that IF-SitePred has a higher average precision (AP) than P2Rank and DeepPocket for multi-structure prediction (Figure 5(b)). While the F1 scores were calculated based on a binary classification with only residues predicted as ligand-binding in at least 3 of 10 structures regarded as positive, the iso-F1 curves applied to the precision-recall curve show that regardless of threshold used, IF-SitePred will achieve a higher F1 score.

**Figure 5:**
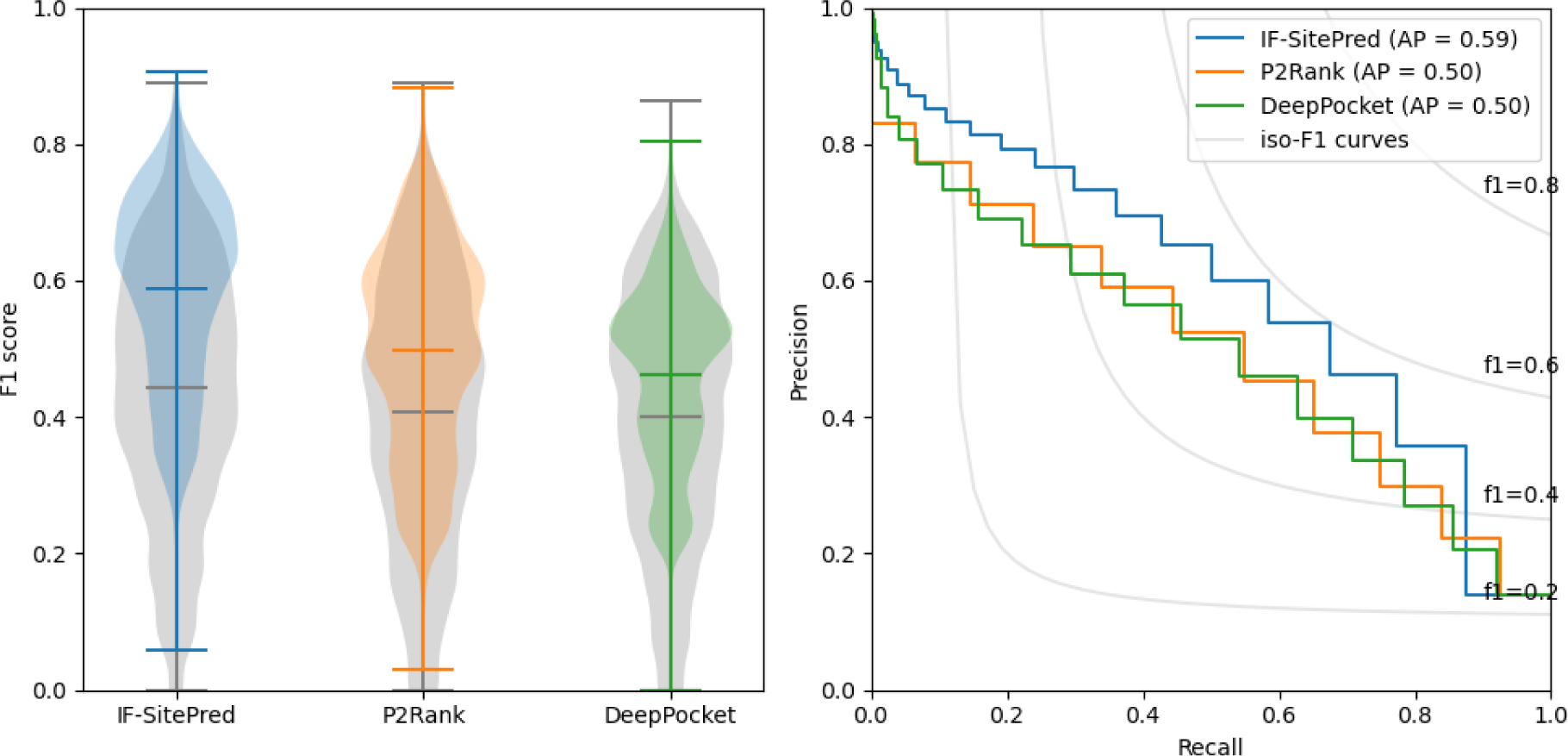
A comparison of the benefits of combining predictions for multiple structures. **Left:** Predictive power for binding residue annotation improves when predictions for 10 structures were combined for all three methods (blue, orange, green) compared to when predictions for single structures were used (grey). IF-SitePred had a slightly higher original F1 score, and also saw the greatest improvement of the three methods, increasing to 0.59. **Right:** Precision-recall curves for all three methods reveal that IF-SitePred has a higher average precision (AP) (0.59) than P2Rank (0.50) and DeepPocket (0.50). Iso-F1 curves are shown in grey, demonstrating that IF-SitePred achieves higher F1 scores across all probability thresholds.

These results suggest that where multiple protein structures are available or can be generated, IF-SitePred is able to take greater advantage of the available data to outperform DeepPocket and P2Rank consistently.

To explore why IF-SitePred was able to take advantage of the information contained in multiple structures better than P2Rank and DeepPocket, we examined the rates of true positives (correctly-labelled ligand-binding residues) and false positives (non-ligand-binding residues labelled as ligand-binding) for residues that were predicted as positive in at least one of the 10 randomly-selected MD structures of the same protein (Figure 6). We found that the number of false positives was reduced in residues that were predicted as positive in more structures, while the number of true positives remained similar. However, IF-SitePred had consistently fewer false positives without a significant reduction in rate of true positives. Conversely, DeepPocket had the highest number of false positives of the three methods, with the number of false positives that were predicted in all 10 structures as ligand-binding almost triple those for P2Rank or IF-SitePred.

**Figure 6:**
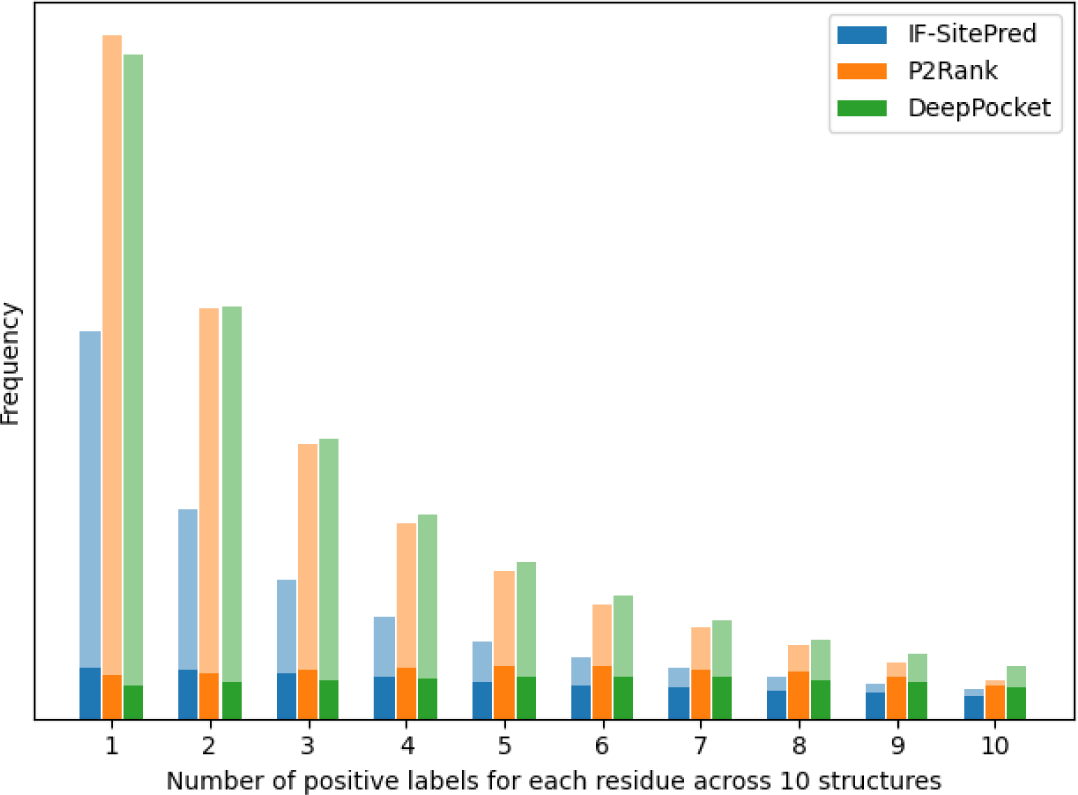
Number of positive labels for each residue of 10 randomly-selected structures. IF-SitePred predicted fewer residues incorrectly as ligand-binding. The frequencies of correctly-labelled (solid colour) and incorrectly-labelled (translucent) residues across 10 randomly-selected MD structures of the same protein are shown. Non-ligand-binding residues were most likely to be incorrectly labelled as ligand-binding in 0 or 1 structure, whereas ligand-binding residues had a roughly equal likelihood of being labelled as ligand-binding up to 10 times. IF-SitePred consistently had a lower rate of labelling non-ligand-binding residues as ligand-binding, with only a small reduction in the number of correctly-labelled ligand-binding residues compared to other methods.

## Discussion

### IF-SitePred binding site prediction is competitive with state-of-the-art methods

We developed IF-SitePred, a binding site prediction method that labels residues based on embeddings generated by ESM-IF1, followed by point cloud clustering to determine the centres of predicted binding sites. We found that IF-SitePred’s performance is comparable with state-of-the-art tools on both PDB protein structures and their equivalent AF2 structures, showing that the ESM-IF1 embeddings contain important information on ligand-binding behaviour of protein residues. When considering the top-3 ranked binding sites on each protein, IF-SitePred achieved the highest success rates of the methods included in this study, detecting at least one correct binding site in around 93% of proteins.

In previous studies where different test sets are used, different methods rank differently: in the DeepPocket analysis [22], FPocket has a much lower success rate than DeepPocket, and DeepPocket significantly outperforms P2Rank. This indicates that even with large test sets, the composition of the set can make a significant difference to results. However, we have shown that in these test sets, IF-SitePred is competitive with state-of-the-art tools on both PDB and AF2 structures, despite no explicit featurisation and a very simplistic pocket ranking strategy.

### IF-SitePred outperforms P2Rank and DeepPocket on low-accuracy predicted structures

When analysing the AF2 structures, we found that the accuracy of the predicted structures was far higher than expected for true novel proteins. We carried out MD simulations on AF2 structures to study at what level of structural error binding site prediction was no longer successful. We found that IF-SitePred, P2Rank and DeepPocket exhibited similar correlations between RMSD of the protein structure (to the PDB structure) and top-1 prediction success: 60% success is achieved on structures with 2Å RMSD or better, while at 4Å RMSD success rate drops to around 40%. However, when considering top-3 prediction success, IF-SitePred is able to maintain success levels around 5% higher than comparators across all RMSD values measured.

While this analysis was performed on a small test set, the protein structures used were more representative of what is expected of novel proteins with no known homologues. Given that IF-SitePred is able to predict binding sites with higher success on these less accurate structures, these results suggest that IF-SitePred is more likely to correctly predict binding sites on the surface of protein structure predictions with low or unknown accuracy.

### IF-SitePred benefits from using combination prediction methods

Proteins are dynamic systems; experimental or computationally derived structures are only snapshots at single points in time, which can limit our prediction success. Additionally, ligand-binding residues are far fewer in number than those that do not bind ligands, which makes it difficult to include all possible data points in the training set for a machine learning model without overfitting on the less represented data. As a first strategy to address these issues, we adopted an ensembling strategy for which we trained multiple models on stratified sub-samples of the dataset, so that no single model was exposed to duplicate data, but in combination, all data was used in the training process. As a second strategy, we used MD simulations and generated additional structures to capture different snapshots of the same protein with the aim to improve overall prediction accuracy.

We found that both of these strategies yield improvements in prediction accuracy. The first strategy improved prediction F1 score from 0.43 to 0.59. Interestingly, for the second strategy, the gain in performance was greatest for IF-SitePred, indicating that IF-SitePred would be able to consistently outperform other methods for binding site prediction where multiple structures of the protein of interest are available, via a reduced rate of false positives without significant loss in recall of true positives.

There are various ways in which multiple structures of the protein of interest may become available. In this paper we used basic MD simulations to generate an ensemble of structures that greatly increase the information available to binding site prediction tools. With the availability of multiple protein structure prediction tools, it is possible to use a variety of tools to generate different structural predictions, which would improve upon the information provided by just one structural prediction. Additionally, several tools have been developed specifically to generate conformational ensembles of protein structures, such as idpGAN [49], which was trained on MD trajectories. Experimentally-determined structures also have the potential to be used as a conformational ensemble, such as where structures have been determined under different conditions.

## Conclusions

As predicted protein structures become the initial input for tools locating ligand binding sites, it is important to evaluate whether these structures are accurate, and develop binding site prediction tools which are robust to errors in structure predictions.

In this study we describe IF-SitePred, a protein binding site prediction tool that is competitive with state-of-the-art tools on high-accuracy AF2 structures. However, we found that the AF2 models used to evaluate binding site prediction methods had far higher accuracy than would be expected for free modelling targets, which would be the primary targets for template-free binding site prediction. Therefore, we examined how binding site prediction tools perform on structures which are representative of novel protein targets.

To evaluate how the inaccuracies in the protein structure prediction impact binding site prediction, we used MD simulations to generate models of the target proteins with varying accuracy and found that IF-SitePred can consistently outperform competitors when predicting three binding site centres on lower-accuracy structures. By taking the most popular predictions on 10 medium-accuracy structures (1-4Å RMSD), predictions made by IF-SitePred can be improved upon significantly, whereas competitors benefit less. This result suggests that by representing the protein as a set of ESM-IF1 embeddings, it is possible to take greater advantage of the diversity in the ensemble of structures compared with the explicit featurisation used for P2Rank and DeepPocket.

Where the accuracy of a predicted structure for a novel target is thought to be poor, our results suggest that IF-SitePred will provide the most reliable binding site prediction of the tools currently available. Additionally, where a group of structures can be generated using MD or by making many structural predictions (by using one or multiple structure prediction tools), the accuracy of binding site prediction can be enhanced further. By accurately locating ligand-binding sites on predicted protein structures, exploration of the target’s function and druggability can begin.

## List of abbreviations

GVP: Geometric Vector Perceptron
PPI: Protein-Protein Interaction
PDB: Protein Data Bank
AF2: AlphaFold2
RMSD: Root Mean Squared Deviation
pLDDT: predicted Local Distance Difference Test
CASP: Critical Assessment of Structure Prediction
FM: Free Modelling
MD: Molecular Dynamics
DBSCAN: Density-Based Spatial Clustering of Applications with Noise
HAP: HOLO4K-AlphaFold2 Paired
HAP-small: HOLO4K-AlphaFold2 Paired (small)
API: Application Programming Interface
GDBT: Gradient Boosted Decision Trees
AUROC: Area Under the Reciever Operating Characteristic curve
DCA: Distance from site Centre to ligand Atom
DCC: Distance from site Centre to ligand Centre
DVO: Discretized Volume Overlap
IoU: Intersection over Union
CNN: Convolutional Neural Network
USRCAT: Ultrafast Shape Recognition with Credo Atom Types
GPCR: G-Protein Coupled Receptor
AP: Average Precision

## Declarations

### Availability of data and materials

Code and trained models are available at https://github.com/oxpig/binding-sites, along with lists of test structures in the HAP, HAP-small and training sets. All training structures are available via the PDB (https://www.rcsb.org/), and all test structures are available from the PDB and from the AlphaFold Protein Structure Database (https://alphafold.ebi.ac.uk/).

### Authors’ contributions

AC developed the methods and wrote the text, with input and support from CMD and RS. All authors reviewed and edited the manuscript.

## Acknowledgements

We extend thanks to Daniel Nissley for his help developing the molecular dynamics simulations.

## Supplementary Information

**Figure S1:**
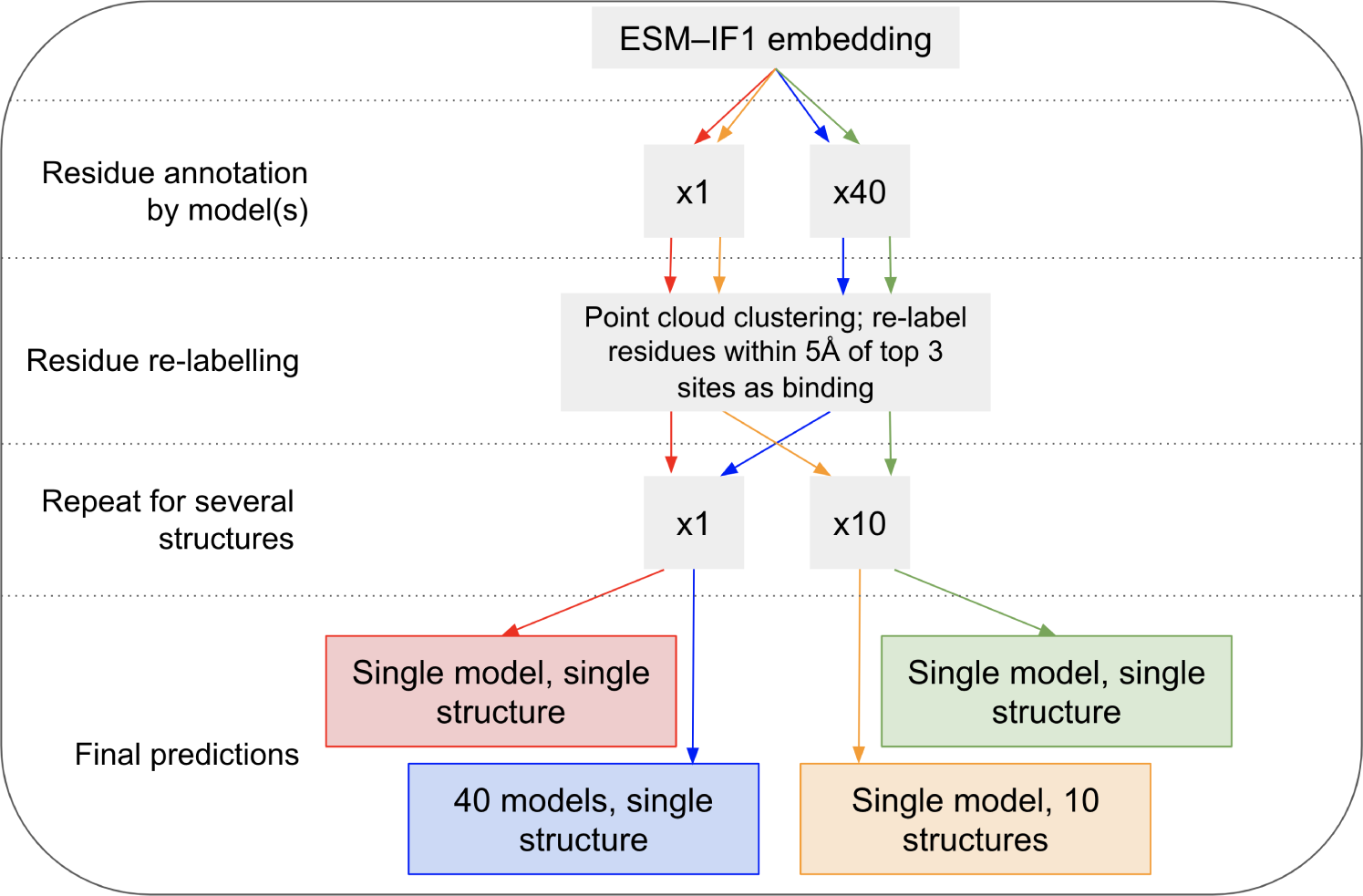
Pathways implemented to compare multiple models on multiple structures. The four pathways implemented to compare the F1 scores achieved when predicting ligand-binding residues on 21 MD structures. These allowed us to explore the benefit gained from using multiple predictive models and from making predictions on multiple MD structures.

**Table S1:**
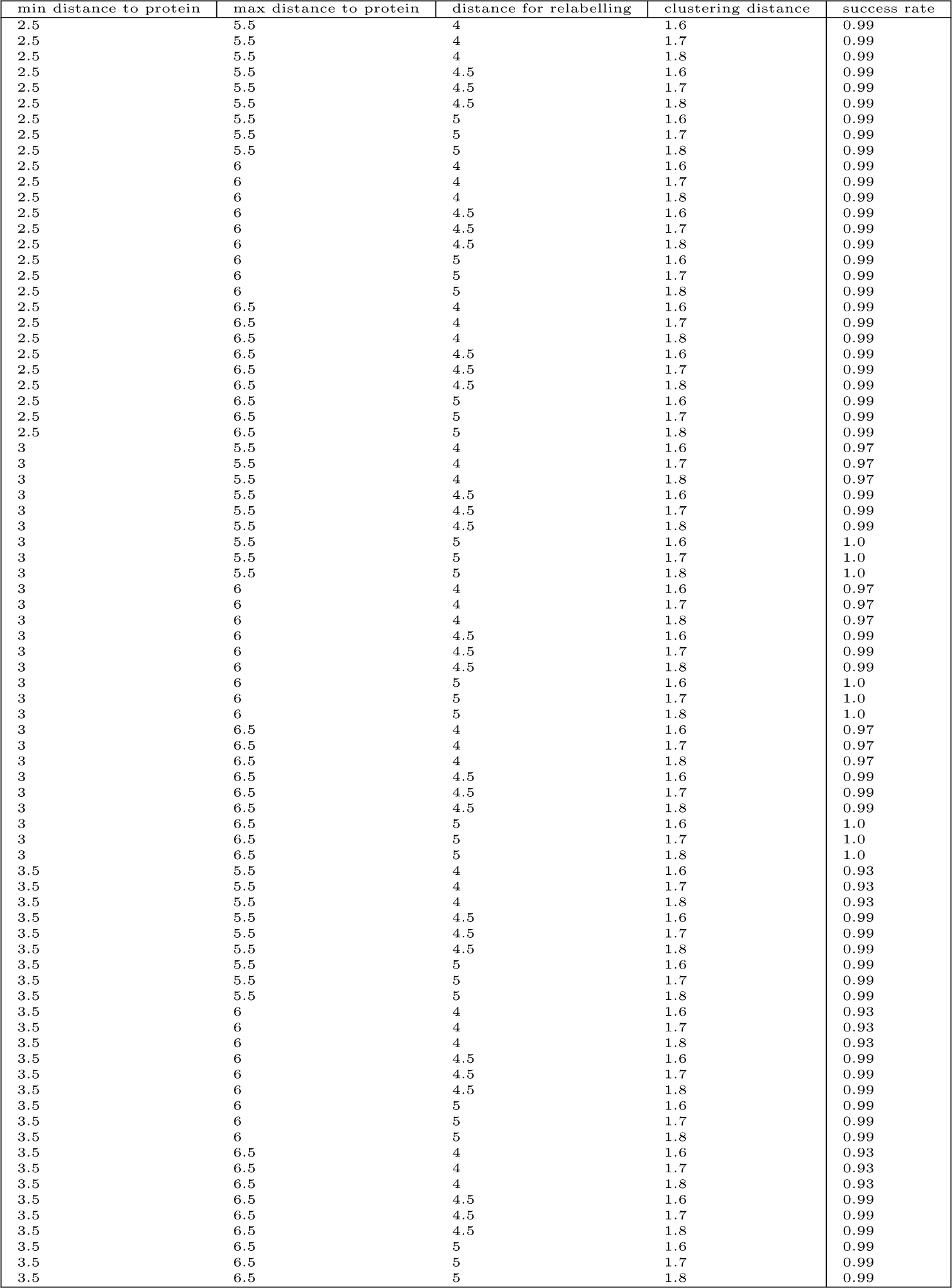
Top-1 success rate on training set using varying distance thresholds. Success rate is largely stable to small perturbations in distance thresholds for point cloud generation and clustering. Thresholds for the following were varied: minimum distance from point to protein; maximum distance from point to protein; maximum distance from ligand-binding point to protein residue for residue to be relabelled as ligand-binding; maximum distance between two points within a cluster. Where the distance for relabelling residues as ligand-binding was lowered (4Å), success rate was reduced as low as 0.93, however all other combinations of thresholds yielded success rates of 0.99-1.00.

