## Supplementary information for "Learnt representations of proteins can be used for accurate prediction of small molecule binding sites on experimentally determined and predicted protein structures"

**Figure S1: Pathways implemented to compare multiple models on multiple structures**

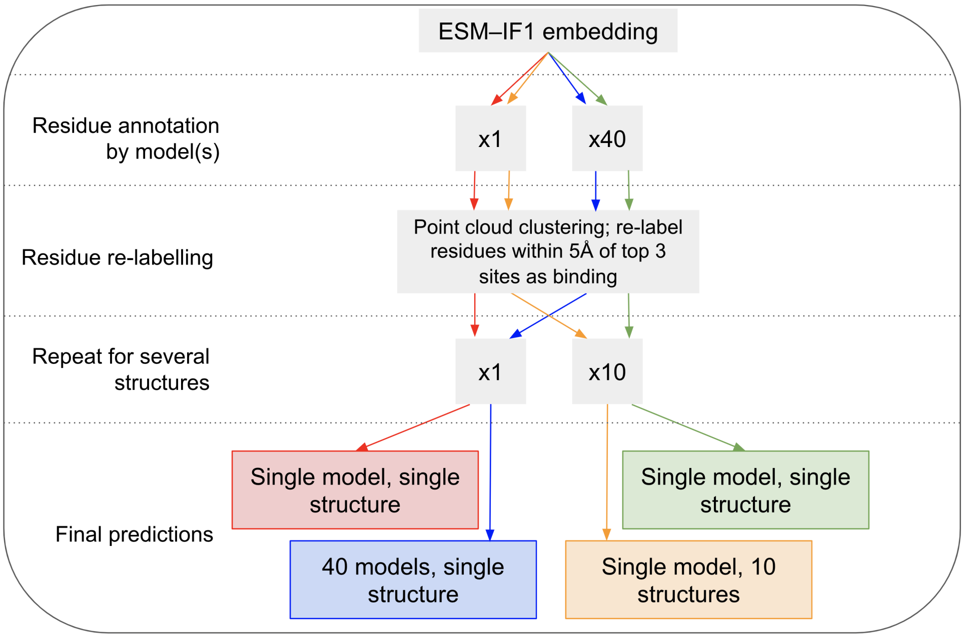


Figure S1: The four pathways implemented to compare the F1 scores achieved when predicting ligand-binding residues on 21 MD structures. These allowed us to explore the benefit gained from using multiple predictive models and from making predictions on multiple MD structures.

### Table S1: Top-1 success rate on training set using varying distance thresholds

| min distance to protein | max distance to protein | distance for relabelling | clustering distance | success rate |
|---|---|---|---|---|
| 2.5 | 5.5 | 4 | 1.6 | 0.99 |
| 2.5 | 5.5 | 4 | 1.7 | 0.99 |
| 2.5 | 5.5 | 4 | 1.8 | 0.99 |
| 2.5 | 5.5 | 4.5 | 1.6 | 0.99 |
| 2.5 | 5.5 | 4.5 | 1.7 | 0.99 |
| 2.5 | 5.5 | 4.5 | 1.8 | 0.99 |
| 2.5 | 5.5 | 5 | 1.6 | 0.99 |
| 2.5 | 5.5 | 5 | 1.7 | 0.99 |
| 2.5 | 5.5 | 5 | 1.8 | 0.99 |
| 2.5 | 6 | 4 | 1.6 | 0.99 |
| 2.5 | 6 | 4 | 1.7 | 0.99 |
| 2.5 | 6 | 4 | 1.8 | 0.99 |
| 2.5 | 6 | 4.5 | 1.6 | 0.99 |
| 2.5 | 6 | 4.5 | 1.7 | 0.99 |
| 2.5 | 6 | 4.5 | 1.8 | 0.99 |
| 2.5 | 6 | 5 | 1.6 | 0.99 |
| 2.5 | 6 | 5 | 1.7 | 0.99 |
| 2.5 | 6 | 5 | 1.8 | 0.99 |
| 2.5 | 6.5 | 4 | 1.6 | 0.99 |
| 2.5 | 6.5 | 4 | 1.7 | 0.99 |
| 2.5 | 6.5 | 4 | 1.8 | 0.99 |
| 2.5 | 6.5 | 4.5 | 1.6 | 0.99 |
| 2.5 | 6.5 | 4.5 | 1.7 | 0.99 |
| 2.5 | 6.5 | 4.5 | 1.8 | 0.99 |
| 2.5 | 6.5 | 5 | 1.6 | 0.99 |
| 2.5 | 6.5 | 5 | 1.7 | 0.99 |
| 2.5 | 6.5 | 5 | 1.8 | 0.99 |
| 3 | 5.5 | 4 | 1.6 | 0.97 |
| 3 | 5.5 | 4 | 1.7 | 0.97 |
| 3 | 5.5 | 4 | 1.8 | 0.97 |
| 3 | 5.5 | 4.5 | 1.6 | 0.99 |
| 3 | 5.5 | 4.5 | 1.7 | 0.99 |
| 3 | 5.5 | 4.5 | 1.8 | 0.99 |
| 3 | 5.5 | 5 | 1.6 | 1.0 |
| 3 | 5.5 | 5 | 1.7 | 1.0 |
| 3 | 5.5 | 5 | 1.8 | 1.0 |
| 3 | 6 | 4 | 1.6 | 0.97 |
| 3 | 6 | 4 | 1.7 | 0.97 |
| 3 | 6 | 4 | 1.8 | 0.97 |
| 3 | 6 | 4.5 | 1.6 | 0.99 |
| 3 | 6 | 4.5 | 1.7 | 0.99 |
| 3 | 6 | 4.5 | 1.8 | 0.99 |
| 3 | 6 | 5 | 1.6 | 1.0 |
| 3 | 6 | 5 | 1.7 | 1.0 |
| 3 | 6 | 5 | 1.8 | 1.0 |
| 3 | 6.5 | 4 | 1.6 | 0.97 |
| 3 | 6.5 | 4 | 1.7 | 0.97 |
| 3 | 6.5 | 4 | 1.8 | 0.97 |
| 3 | 6.5 | 4.5 | 1.6 | 0.99 |
| 3 | 6.5 | 4.5 | 1.7 | 0.99 |
| 3 | 6.5 | 4.5 | 1.8 | 0.99 |
| 3 | 6.5 | 5 | 1.6 | 1.0 |
| 3 | 6.5 | 5 | 1.7 | 1.0 |
| 3 | 6.5 | 5 | 1.8 | 1.0 |
| 3.5 | 5.5 | 4 | 1.6 | 0.93 |
| 3.5 | 5.5 | 4 | 1.7 | 0.93 |
| 3.5 | 5.5 | 4 | 1.8 | 0.93 |
| 3.5 | 5.5 | 4.5 | 1.6 | 0.99 |
| 3.5 | 5.5 | 4.5 | 1.7 | 0.99 |
| 3.5 | 5.5 | 4.5 | 1.8 | 0.99 |
| 3.5 | 5.5 | 5 | 1.6 | 0.99 |
| 3.5 | 5.5 | 5 | 1.7 | 0.99 |
| 3.5 | 5.5 | 5 | 1.8 | 0.99 |
| 3.5 | 6 | 4 | 1.6 | 0.93 |
| 3.5 | 6 | 4 | 1.7 | 0.93 |
| 3.5 | 6 | 4 | 1.8 | 0.93 |
| 3.5 | 6 | 4.5 | 1.6 | 0.99 |
| 3.5 | 6 | 4.5 | 1.7 | 0.99 |
| 3.5 | 6 | 4.5 | 1.8 | 0.99 |
| 3.5 | 6 | 5 | 1.6 | 0.99 |
| 3.5 | 6 | 5 | 1.7 | 0.99 |
| 3.5 | 6 | 5 | 1.8 | 0.99 |
| 3.5 | 6.5 | 4 | 1.6 | 0.93 |
| 3.5 | 6.5 | 4 | 1.7 | 0.93 |
| 3.5 | 6.5 | 4 | 1.8 | 0.93 |
| 3.5 | 6.5 | 4.5 | 1.6 | 0.99 |
| 3.5 | 6.5 | 4.5 | 1.7 | 0.99 |
| 3.5 | 6.5 | 4.5 | 1.8 | 0.99 |
| 3.5 | 6.5 | 5 | 1.6 | 0.99 |
| 3.5 | 6.5 | 5 | 1.7 | 0.99 |
| 3.5 | 6.5 | 5 | 1.8 | 0.99 |

Table S1: Success rate is largely stable to small perturbations in distance thresholds for point cloud generation and clustering. Thresholds for the following were varied: minimum distance from point to protein; maximum distance from point to protein; maximum distance from ligand-binding point to protein residue for residue to be relabelled as ligand-binding; maximum distance between two points within a cluster. Where the distance for relabelling residues as ligand-binding was lowered (4Å), success rate was reduced as low as 0.93, however all other combinations of thresholds yielded success rates of 0.99-1.00.
